# The antibody-based delivery of interleukin-12 to solid tumors boosts NK and CD8^+^ T cell activity and synergizes with immune check-point inhibitors

**DOI:** 10.1101/684100

**Authors:** Emanuele Puca, Philipp Probst, Marco Stringhini, Patrizia Murer, Giovanni Pellegrini, Samuele Cazzamalli, Cornelia Hutmacher, Baptiste Gouyou, Sarah Wulhfard, Mattia Matasci, Alessandra Villa, Dario Neri

## Abstract

We describe the cloning and characterization of a novel fusion protein (termed L19-mIL12), consisting of murine interleukin-12 in single-chain format, sequentially fused to the L19 antibody in tandem diabody format. The fusion protein bound avidly to the cognate antigen (the alternatively-spliced EDB domain of fibronectin), retained the activity of the parental cytokine and was able to selectively localize to murine tumors *in vivo*, as shown by quantitative biodistribution analysis. L19-mIL12 exhibited a potent anti-tumor activity in immunocompetent mice bearing CT26 carcinomas and WEHI-164 sarcomas, which could be boosted by combination with check-point blockade, leading to durable cancer eradication. L19-mIL12 also inhibited tumor growth in mice with Lewis lung carcinoma (LLC), but in this case cancer cures could not be obtained, both in monotherapy and in combination. A microscopic analysis and a depletion experiment of tumor-infiltrating leukocytes illustrated the contribution of NK cells and CD8^+^ T cells for the anti-cancer activity observed in both tumor models. Upon L19-mIL12 treatment, the density of regulatory T cells (Tregs) was strongly increased in LLC, but not in CT26 tumors. A FACS analysis also revealed that the majority of CD8^+^ T cells in CT26 tumors were specific to the retroviral AH1 antigen.

**NOVELTY AND IMPACT STATEMENT:** In this study, we describe the generation of a novel fusion protein consisting of murine inter-leukin-12 fused to the L19 antibody (specific to the EDB domain of fibronectin). L19-mIL12 revealed favourable tumor to organ ratios 24h after intravenous administration and it was able to cure 60% of CT26 tumor-bearing mice. From a clinical perspective, the rapid clearance from circulation should ease the administration to patients as infusions could be stopped upon onset of side-effects.

## INTRODUCTION

Immune check-point inhibitors (such as antibodies targeting PD-1, PD-L1 or CTLA-4) are revolutionizing the treatment of patients with various types of malignancies ^1–3^. These biopharmaceuticals can be considered as “foundational drugs”, to which other therapeutic regimens may be added, in order to achieve optimal anti-cancer activity ^4, 5^. In particular, inhibition of the PD-1/PD-L1 axis may help reverse an exhausted phenotype in CD8^+^ T cells, which have previously encountered tumor-associated antigens ^6–8^. As a consequence, these therapeutic strategies are likely to be more effective, when a substantial number of tumor-specific lymphocytes are present within the solid tumor mass ^9^.

Cytokines are a class of immunomodulatory proteins, which have been considered as biopharmaceuticals for cancer therapy ^10, 11^. In particular, certain pro-inflammatory cytokines (such as interleukin-2, −12 and −15) could potentially be used to boost the activity of T cells and NK cells within the neoplastic mass. However, most cytokine products cause side effects at low doses (sometimes even at 1μg/Kg in patients), thus preventing dose escalation to therapeutically active regimens ^12^. The fusion of cytokines to antibodies, capable of selective localization to the tumor mass, has been proposed as a means to increase the therapeutic index of these biopharmaceuticals ^13–15^. Some antibody-cytokine fusions are currently being investigated in the clinic for the treatment of different types of malignancies ^16–24^.

Interleukin-12 (IL12) is a heterodimeric cytokine consisting of the p40 and p35 subunits, linked by a disulfide bond ^25, 26^. The protein is predominantly produced by Antigen Presenting Cells (APCs) upon recognition of “danger signals” such as pathogen associated molecular patterns (PAMPs) ^25, 27^. IL12 can bind to its cognate receptor, composed of IL12R*β*1 and IL12R*β*2, driving the activation of interferon-*γ* (IFN-*γ*) transcription factors through a Jak/STAT pathway. IFN-*γ* is one of the most relevant mediators of the pro-inflammatory effect induced by of IL12 and favours a T_H_1 differentiation of CD4^+^ T-cells. Moreover, IL12 has also been shown to (i) increase the proliferation of NK, CD4^+^ and CD8^+^ T cells, (ii) increase antigen processing and presentation by APCs, (iii) augment ADCC against cancer cells, (iv) inhibit Treg activity, promoting their polarization towards a T_H_1 phenotype, (v) enhance IgG production and (vi) alter expression patterns of endothelial adhesion molecules such as VCAM-1 ^27^.

For the last two decades, various groups, including our own, have worked on the engineering of antibody-IL12 fusions, since the heterodimeric nature of IL12 poses various opportunities for protein engineering, leading to formats with different pharmacokinetic and pharmacodynamic properties ^28–30^. Two antibody-IL12 fusions have recently entered clinical trials for treatment of patients with cancer: BC1-IL12 ^16^ and NHS-IL12 (NCT01417546) ^31^. Both fusion proteins are based on intact immunoglobulin structures. However, antibody fragments may be preferable for the generation of IL12 fusion proteins, in order to achieve a more rapid clearance from circulation and better tumor:organ ratios ^32^.

Here, we describe a novel fusion protein (termed L19-mIL12), consisting of a single-chain murine interleukin-12, sequentially fused to the L19 antibody in tandem diabody format. L19 specifically recognizes the alternatively-spliced EDB domain of fibronectin, a highly selective tumor-associated antigen. Over the last 20 years, several publications (based on the L19 antibody or similar phage-derived antibodies) have confirmed EDB as a pan-tumoral antigen, which is undetectable in most normal tissues, exception made for certain structures of the female reproductive system ^33^. L19 recognizes both murine and human EDB with identical affinity, since the domain sequence is completely conserved between the two species ^34^. However, since the human isoform of IL12 does not cross react with the cognate murine receptor, preclinical studies in mice were performed using the murine IL12 as therapeutic payload ^35^.

The L19-mIL12 product, which retained full activity in terms of both antigen binding and cytokine activity, was potently active against CT26 carcinomas and cured all mice when combined with CTLA-4 or PD-1 blockade. The fusion protein was also potently active also against LLC but, in this case, complete tumor eradications could not be achieved, not even in combination with immune check-point inhibition. While in both models L19-mIL12 mediated an increased density of intratumoral CD8^+^ T cells and NK cells, Treg density was dramatically increased only in LLC tumors. Staining of CD8^+^ T cells with tetramer reagents in the CT26 model revealed a striking difference between tumor-infiltrating lymphocytes and their counterparts in tumor-draining lymph nodes. The results of our study suggest that a fully human L19-IL12 fusion protein may be considered for clinical development activities.

## MATERIALS AND METHODS

### Cell lines, animals and tumor models

CHO-S (Invitrogen; CVCL_7183), CT26.WT (ATCC; CVCL_7256), LL/2 (ATCC; CVCL_4358), F9 (ATCC; CVCL_0259) and WEHI-164 (ATCC; CVCL_2251) were expanded and stored as cryopreserved aliquots in liquid nitrogen. Cells were grown according to the manufacturer’s protocol and kept in culture for no longer than 14 passages. Authentication of the cell lines including post-freeze stability, growth properties and morphology, test for mycoplasma contamination, isoenzyme assay, and sterility were performed by the cell bank before shipment. All experiments were performed with mycoplasma-free cells.

Seven to eight-week-old female BALB/c, C57BL/6 and 129/svEv mice were obtained from Janvier. 4*10^6^ cells (CT26), 3*10^6^ cells (WEHI-164), 2*10^6^ cells (LLC) and 10*10^6^ cells (F9) were implanted subcutaneously in the left flank of the mice. Experiments were performed under a project license (license number 27/2015, 04/2018) granted by the Veterinäramt des Kantons Zürich, Switzerland, in compliance with the Swiss Animal Protection Act (TSchG) and the Swiss Animal Protection Ordinance (TSchV).

### Cloning, expression and *in vitro* protein characterization

The format chosen for L19-mIL12 was inspired by previous work in our laboratory with F8 antibody derivatives ^30^. The sequence of the gene is reported in **Supplementary Figure 1**. The insert was cloned into NheI/NotI of pcDNA3.1 (+) (Invitrogen), allowing the expression in mammalian cells. L19-mIL12 was expressed using transient gene expression in Chinese Hamster Ovary (CHO) cells, using procedures previously described for other antibody fusions ^36, 37^. The product was purified from the cell culture medium by affinity chromatography using a Protein A affinity column and analyzed by SDS-PAGE, Size Exclusion Chromatography (Superdex200 10/300GL, Healthcare) and Surface Plasmon Resonance analysis (Biacore S200, GE Healthcare) on an EDB antigen-coated sensor chip (CM5, GE Healthcare).

### Bioactivity assay

L19-mIL12, KSF-mIL12 and recombinant mIL12 (BioLegend) were subjected to a splenocyte IFN-*γ* release assay. Splenocytes were isolated from freshly dissected spleens of BALB/c mice. After red blood cell lysis, splenocytes were resuspended at 5*10^6^ cells/mL in RPMI-1640 (Thermo Fisher) supplemented with antibiotic-antimycotic (Thermo Fisher; 15240-062) and 10% Fetal Bovine Serum (Gibco; 10270-106). 100*μ*L of the cell suspension was incubated for 5 days at 37°C and 5% CO_2_ with a serial dilution of the IL12 derivatives. Cultured supernatants were analysed by a sandwich enzyme-linked immunosorbent assay (ELISA) using 5 *μ*g/mL of the monoclonal anti-mouse IFN-*γ* antibody (eBioscience) for capture and 1 *μ*g/mL of the polyclonal biotinylated anti-mouse IFN-*γ* antibody (PeproTech) for detection.

### Cytokine analysis in tumor extracts and plasma

For a quantitative analysis of cytokines in tumors extracts and in plasma, CT26 and LLC bearing mice were euthanized 24h after 3 intravenous injections, every third day, of PBS or 12*μ*g of L19-mIL12. BALB/c mice were immediately exsanguinated and the blood was left to clot for 1h at room temperature in Microtainer tubes containing lithium heparin (BD Microtainer Tube). Plasma was collected after centrifugation, aliquoted and frozen at −80 °C until cytokine assays were performed.

Excised Lewis Lung and CT26 carcinomas were suspended in 2mL of Tris-HCl buffer (50mM, pH 7.4) containing NaCl (0.6M), Triton X-100 (0.2%), BSA (0.5%) and freshly dissolved protease inhibitors (1mM benzamidine, 0.1mM benzethonium chloride and 0.1mM phenylmethylsulfonyl fluoride). Tumors were homogenized with QIAGEN Tissue Lyser (4 × 1’, 30 Hz) using a 5mm stainless steel bead (Qiagen, Hombrechtikon, Switzerland) and sonicated for 20s. Supernatants were collected after centrifugation, aliquoted and frozen at −80 °C until cytokine assays were performed.

Cytokine levels were eventually measured using a multiplexed particle-based flow cytometric assay ^38^. The kits Luminex Screening assay (LXSAMS-5 and LXSAMS-6) and Luminex Perfomance Assay (LTGM100) were purchased from Bio-Techne (Zug, Switzerland). The procedure closely followed the manufacturer’s instructions and the analysis was conducted using a conventional flow cytometer (Guava EasyCyte 8HT, Millipore, Zug, Switzerland) at the Cytolab facility (Ostring 48, CH-8105 Regensdorf).

### Immunofluorescence studies

The expression of EDB (+)-fibronectin in CT26, LLC, MC38, F9 and WEHI-164 tumors was studied on ice-cold acetone-fixed 8-10*μ*m cryostat sections stained with FITC labeled IgG(L19) (FITC_F7250 Sigma) (final concentration 2*μ*g/mL) and detected with a Rabbit anti-FITC (Bio-Rad; 4510-7804) and Goat anti-Rabbit AlexaFluor488 (Invitrogen; A1108). For vascular staining, goat anti-CD31 (R&D System; AF3628) and anti-goat AlexaFluor594 (Invitrogen; A11058) antibodies were used.

For *ex vivo* immunofluorescence analysis, mice were injected according to the therapy schedule and euthanized 24h after the last injection. Tumors were excised and embedded in cryoembedding medium (ThermoScientific) and the corresponding cryostat tissue sections (8-10*μ*m thickness) were stained using the following primary antibodies: goat anti-CD31 (R&D System; AF3628), rabbit anti-caspase-3 (Sigma; C8487), rabbit anti-Foxp3 (Invitrogen; 7000914), rabbit anti-Asialo GM1 (Wako; 986-10001), rabbit anti-CD4 (Sino Biological; 50134-R001), rabbit anti-CD8 (Sino Biological; 50389-R208). Primary antibodies were detected with Donkey anti-rabbit AlexaFluor488 (Invitrogen; A11008) and Donkey anti-goat AlexaFluor594 (Invitrogen; A21209). Slides were mounted with fluorescent mounting medium (Dako Agilent) and analyzed with wide field Axioskop2 mot plus microscope (Zeiss) using the AxioVision software (4.7.2 Release, Zeiss). Quantification of the infiltrate analysis, using Image J software, is depicted in **Supplementary Figure 6**. The analysis of infiltrative NK cells stained with the anti-NKp46 antibody (BioLegend; 137602) is shown in **Supplementary Figure 4**.

### *In vivo* quantitative biodistribution analysis

The tumor homing ability of L19-mIL12 was assessed by *in vivo* quantitative biodistribution studies, according to a previously described procedure ^39^. Briefly, 8*μ*g of radioiodinated L19-mIL12 were injected into the lateral tail vein of F9 tumor-bearing 129/svEv mice (Janvier) when tumors had reached approximately 150mm^3^. Mice were sacrificed 24h after the injection and the organs were excised and weighed. Radioactivity of tumors and organs was measured using a Cobra *γ* counter (Packard, Meriden, CT, USA) and expressed as percentage of injected dose per gram of tissue (%ID/g ± SEM; n=2mice).

### Therapy studies and *in vivo* lymphocyte depletion

Mice were monitored daily. Tumor volume was measured using a caliper (volume=length *width*0.52). Mice were intravenously injected with 8.75*μ*g or 12*μ*g of L19-mIL12, starting when tumors reached approximately 100mm^3^, every third day for three times. The therapeutic agent was diluted in Phosphate-Buffer Saline (PBS; Gibco). In the combination groups, mice received L19-mIL12 followed by 200*μ*g of a check-point inhibitor: either αPD-1 (clone 29F.1A12) or aPD-L1 (clone 10F.9G2) or aCTLA-4 (clone 9D9) antibody. All the checkpoint inhibitor antibodies were purchased from BioXCell. Animals were euthanized when tumors reached a maximum of 1500mm^3^. (n=4-5 mice per group)

Cured mice were rechallenged by subcutaneous injection of 4*10^6^ CT26 Colon carcinoma cells on day 49 after the first tumor implantation.

For the *in vivo* depletion of NK cells, CD4^+^ and CD8^+^ T cells, LLC (n=4-5 mice per group) and CT26 Colon carcinoma (n=5 mice per group) bearing mice were injected 3 to 4 times intraperitoneally with 30*μ*g rabbit anti-Asialo GM1 (Wako Chemicals), 250*μ*g rat anti-CD8 (YTS169.4, BioXCell) and 250*μ*g rat anti-CD4 (GK1.5, BioXCell). In the Lewis Lung model, an additional group (n=4) was injected with 250*μ*g anti-CD25 antibody (PC-61.5.3, BioXCell). One saline group (n=5) and one treatment group (n=4-5) were included in both models as controls. The injection of the depleting antibodies was started 2 days prior to therapy initiation and performed every third day for 4 times (except made for the anti-Asialo GM1 that was given 3 times). Animals were euthanized when tumors reached 1500mm^3^ or weight loss exceeded 15%.

### Flow cytometric analysis of CD4^+^ T cells, CD8^+^ T cells and CD8 AH1-specific T cells in tumor draining lymph nodes and in the tumor

CT26 tumor-bearing mice were sacrificed after 1 and 3 injections of PBS or 12*μ*g of L19-mIL12.

Excised tumors were cut in small fragments, passed through a 40*μ*m nylon cell-strainer (EASYstrainer, greiner bio-one), and digested for 2h at 37 at 110rpm in RPMI-1640 (Thermo Fisher, with L-glutamine) containing antibiotic-antimycotic solution, 1mg/mL collagenase II (Thermo Fisher) and 0.1mg/mL DNase I (Roche). Cell suspensions were passed through a 70*μ*m nylon cell-strainer (EASYstrainer, greiner bio-one), repeatedly washed and immediately used for flow cytometry analysis.

Excised tumor draining lymph nodes (dLN) were minced in PBS, treated with Red-Blood Cell Lysis Buffer (BioLegend) and passed through a 40*μ*m nylon cell-strainer (EASYstrainer, greiner bio-one). The cell suspensions were then repeatedly washed and immediately used for flow cytometry analysis.

For flow cytometry analysis, approximately 1 * 10^6^ cells from dLN and 5 × 10^6^ tumor cells were first stained with Zombie Red Fixable Viability Kit (BioLegend) for 30’ at room temperature and then incubated on ice for 15’ with an anti-mouse CD16/32 antibody (BioLegend). Cell surface markers were stained using APC-coupled AH1 tetramers and fluorochrome-conjugated antibodies against CD8a (FITC), CD4 (PerCP), CD44 (APC/Cy7), CD62L (BV421), CD90.2 (PE), PD-1 (BV605) and Tim-3 (BV421) which were all purchased from BioLegend for 1h on ice. The conjugated antibodies were diluted in PBS containing 0.5% bovine serum albumin and 2mM EDTA. For the staining of intracellular markers, cells were permeabilized with Fixation/Permeabilization solution (Fixation/Permeabilization Diluent + Fixation/Permeabilization Concentrate, Invitrogen) for 1h on ice. Afterwards, tumors and dLN were centrifuged and resuspended in Permeabilization Buffer (Invitrogen) with the fluorochrome-conjugated antibodies against Ki67 (Alexa-700, BioLegend) and Foxp3 (BV421, BioLegend) for 1h at room temperature. Cells were analysed on a CytoFLEX cytometer (Beckman Coulter) and data was processed using FlowJo (v.10. Tree Star).

### Histopathological evaluation of BALB/c mice

Healthy BALB/c were euthanized 24h after the third administration of saline or 12μg L19-mIL12. The same treatment schedule of the tumor therapies depicted in **Figure 3** was used. A full necropsy was conducted in each mouse, main organs were removed and fixed in 4% neutral-buffered formalin (Formafix, Hittnau, Switzerland) for 48 h. After fixation tissues were trimmed, dehydrated in graded alcohol and routinely paraffin wax embedded. Consecutive sections (3–5μm thick) were prepared, mounted on glass slides and routinely stained with haematoxylin and eosin (HE).

### Data availability

The data will be made available upon reasonable request.

## RESULTS

### Cloning and characterization of the L19-mIL12 fusion protein

Figure 1A **and** B depict the structure of the vector for expression in mammalian cells and the domain assembly of L19-mIL12. The p40 and p35 subunits of murine IL12 were sequentially fused to the L19 antibody in tandem diabody format ^30^. The gene sequence and full amino acid sequence of the fusion protein are reported in **Supplementary Figure 1**. The product could be purified by Protein A chromatography [**Figure 1C, D**], since the VH domain of the L19 antibody retains Protein A affinity ^34^. L19-mIL12 exhibited a high affinity to recombinant EDB domain of fibronectin in Biacore binding studies [**Figure 1E**]. The ability to induce IFN-*γ* production in BALB/c splenocytes was similar to the one of recombinant murine IL12 and KSF-mIL12 [**Figure 1F**]. When injected 3 times in BALB/c mice at a dose of 12*μ*g, L19-mIL12 mediated an elevation in the levels of IL6, IL10, TNF and IFN-*γ*, while the impact on other cytokines (e.g., IL2, IL4, IL17A) was less pronounced [**Figure 1G, H**]. Injections of L19-mIL12 in BALB/c mice at doses up to 12*μ*g were well tolerated, as revealed by a detailed histopathological analysis [**Supplementary Figure 2**].

**Figure 1.**
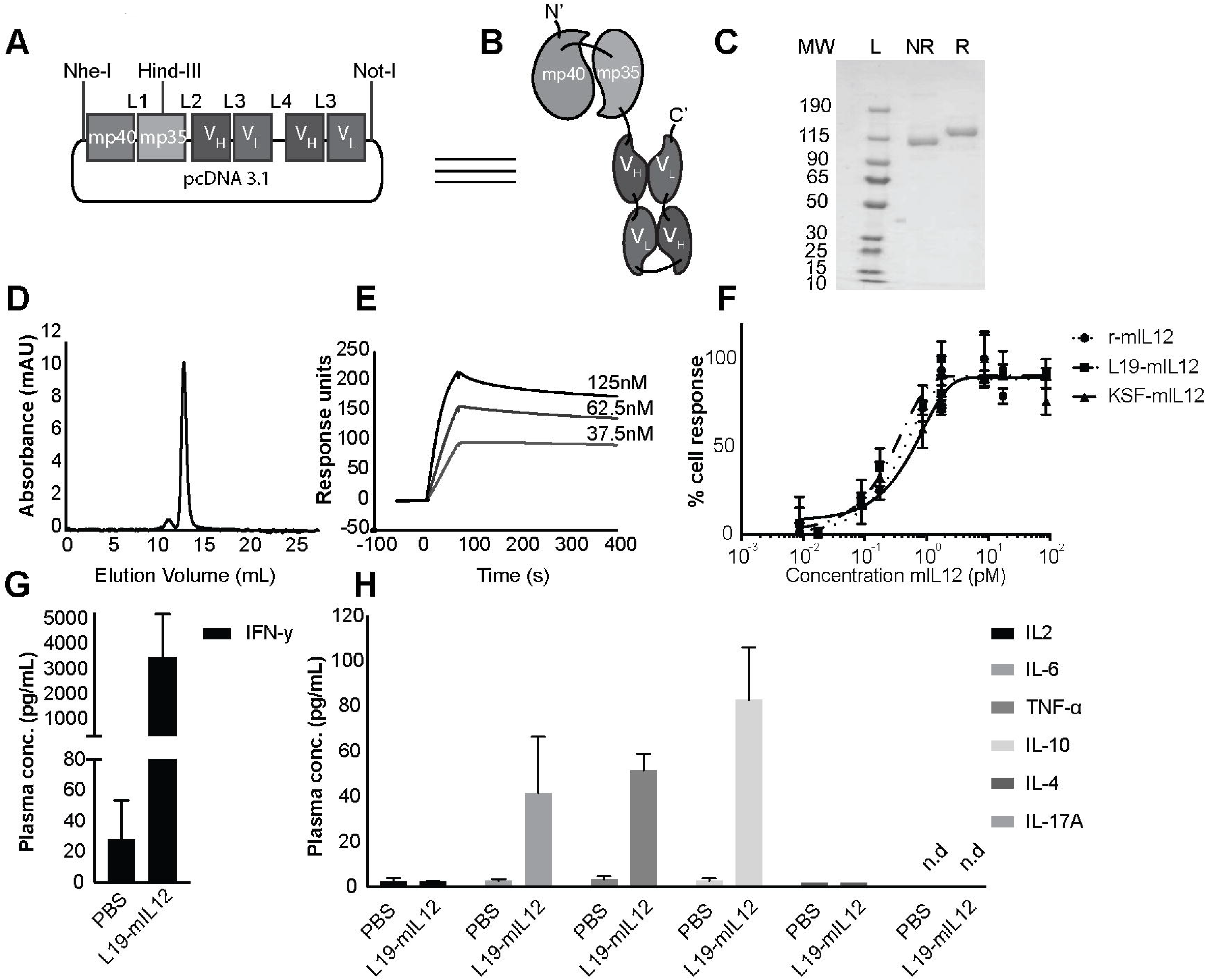
*In vitro* and *in vivo* characterization of L19-mIL12. (**A**) Scheme of the L19-mIL12 construct inserted in the mammalian expression vector pcDNA3.1(+). L1, L2, L3, L4 correspond to Linker1, Linker2, Linker3, Linker4, respectively. (**B**) Scheme of the heterodimeric murine IL12 fused to the L19 antibody in tandem diabody format. (**C**) SDS-PAGE analysis of purified L19-mIL12 under non-reducing (NR) and reducing (R) conditions. L=Ladder. MW=Molecular Weight (**D**) Size-Exclusion Chromatography profile of L19-mIL12. (**E**) Surface Plasmon Resonance analysis of L19-mIL12 on EDB antigen-coated sensor chip. (**F**) IFN-*γ* induction assay by L19-mIL12, KSF-mIL12 and recombinant mIL12 in BALB/c splenocytes. (**G, H**) Plasma concentration of cytokines in BALB/c mice injected every third day for 3 times with 12μg L19-mIL12. Blood samples were collected 24h after the last injection. n.d.=non-detectable.

### Microscopic analysis of the EDB and *in vivo* quantitative biodistribution study

The staining characteristics of the L19 antibody against various types of murine tumors is depicted in **Figure 2A**. While certain tumors (e.g. CT26, LLC) exhibited a predominantly stromal pattern of staining, F9 teratocarcinomas showed a more perivascular expression pattern. Similar findings have been reported for human tumors ^40^. When injected intravenously into immunocompetent mice bearing F9 teratocarcinomas, a radioiodinated preparation of L19-mIL12 revealed a preferential uptake in the solid tumor mass 24 hours after bolus injection, with 7 percent injected dose per gram (%ID/g) of tumor, compared to less than 1% ID/g in blood [**Figure 2B**].

**Figure 2.**
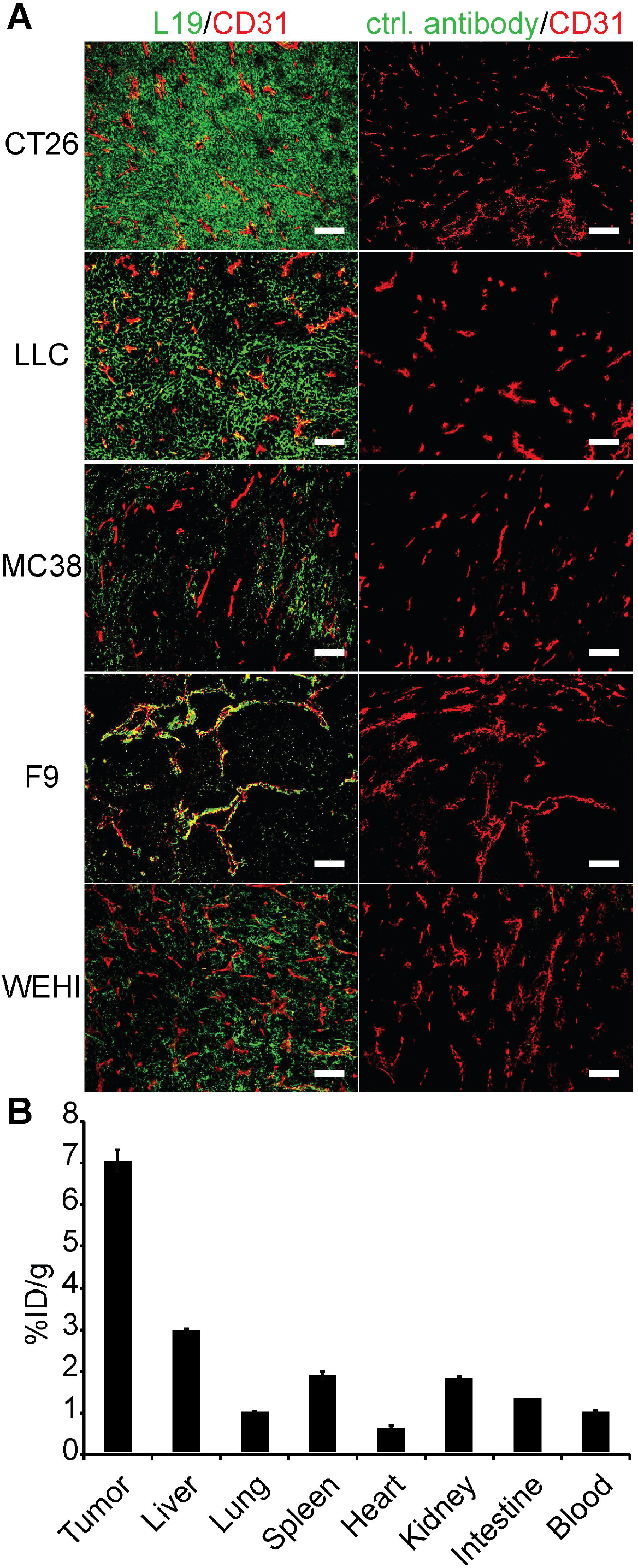
Microscopic analysis of the EDB domain of fibronectin and tumor-targeting study of L19-mIL12. (**A**) Microscopic fluorescence analysis of EDB expression (green) on different tumor mouse models using a FITC-labeled L19 IgG1 or FITC-labeled KSF IgG1 (negative control antibody, specific for an irrelevant antigen). An anti-CD31 antibody was used to stain blood vessels (red). Scale bar=100μm. (**B**) Quantitative biodistribution experiment of 8μg radioiodinated L19-mIL12 24h after intraveneous injection in immunocompetent F9 teratocarcinoma bearing mice. Results are expressed as percentage of injected dose per gram of tissue (%ID/g ±SEM; n=2).

### Therapy experiments

In a first therapy study conducted in BALB/c mice bearing CT26 tumors, L19-mIL12 exhibited a potent tumor-growth inhibition activity at a dose of 8.75μg. The product was very well tolerated, as evidenced by no loss of body weight, and more efficacious than KSF-mIL12 (a fusion protein of irrelevant specificity, directed against hen egg lysozyme and used as a negative control) [**Figure 3A**].

**Figure 3.**
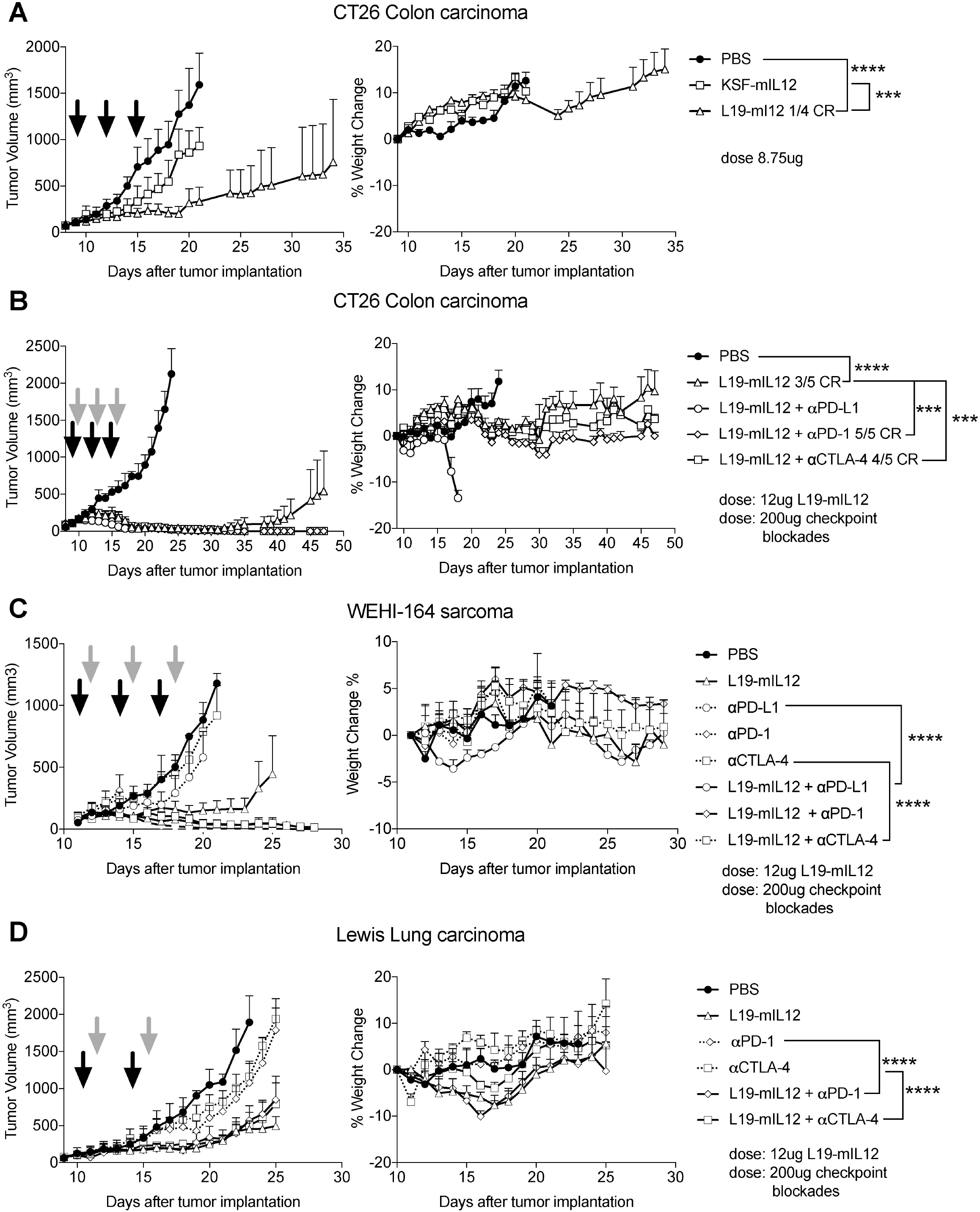
Therapeutic performance of L19-mIL12 against CT26 Colon carcinoma and Lewis Lung carcinoma. (**A**) Therapy study in BALB/c mice bearing CT26 lesions (left) with corresponding body weight change (right). Treatment started on day 9 after cancer cell implantation when tumors reached 110mm^3^. Mice received either PBS, 8.75*μ*g L19-mIL12 or 8.75*μ*g KSF-mIL12 (black arrows) 3 times every 72h (n=4 mice per group). (**B**) Therapy in BALB/c mice bearing CT26 lesions (left) with corresponding body weight change (right). Treatment started on day 9 after cancer cell injection when tumors reached 110mm^3^. Mice received PBS or 12*μ*g of L19-mIL12 (black arrows) alone or in combination with 200*μ*g of a check-point inhibitor (grey arrows) 3 times (n=5 mice per group). A mouse in the combination group of L19-mIL12 + *α*CTLA-4 had to be euthanized on day 17 because of tumor ulceration. (**C**) Therapy in BALB/c mice bearing WEHI-164 lesions (left) with corresponding body weight change (right). The same treatment schedule of (**B**) was used. Mice treated with *α*PD-L1 alone and in combination received only 2 rounds of injections. (n=4-5 mice per group). (**D**) Therapy in C57BL/6 mice bearing LLC lesions with corresponding body weight change (right). Treatment started on day 10 after cancer cell injection when tumors reached 110mm^3^ (n=4 mice per group). Mice received PBS, 200 *μ*g *α*PD-1, 200 *μ*g *α*CTLA-4 or 12*μ*g of L19-mIL12 (black arrows) alone or in combination with 200*μ*g of a check-point inhibitor (grey arrows) twice (n=4 mice per group). Statistical differences were assessed between treatment and control groups. ***, p<0.001; ****, p<0.0001 (regular two-way ANOVA test with Bonferroni post-test). Data represent mean tumor volume and body weight change (±SEM). CR=Complete response.

The preliminary study provided a motivation to increase the dose of L19-mIL12 to 12μg and to test the product in combination with mouse-specific immune check-point inhibitors. In this setting, 3/5 mice could be cured by the L19-mIL12 monotherapy, while 4/5 and 5/5 complete responses could be achieved in combination with CTLA-4 or PD-1 blockade respectively [**Figure 3B**]. The immune check-point inhibitors only showed only minimal single agent activity [**Supplementary Figure 5**]. Surprisingly, the combination with the anti-PD-L1 antibody was not well tolerated and mice had to be prematurely sacrificed, due to the loss of body weight [**Figure 3B**]. In order to minimize PD-L1-related toxicity, mice with WEHI-164 sarcoma were given only two injections of anti-PD-L1 antibody in all study groups [**Figure 3C**]. L19-mIL12 was potently active in inhibiting tumor growth also in this model, leading to durable complete responses in all combination groups. In the monotherapy group, only one mouse enjoyed a durable complete response with L19-mIL12, whereas tumors progressed upon treatment with immune checkpoint inhibitors alone [**Figure 3C**].

A similar investigation was conducted in C57BL/6 mice, bearing subcutaneous LLC tumors. In this case, a tumor growth inhibition activity of L19-mIL12 could be observed, but mice could not be cured, not even in combination with immune check-point inhibitors [**Figure 3D**].

### Microscopic analysis of tumor-infiltrating lymphocytes

In order to gain more insights into the different performance of L19-mIL12 in CT26 and LLC tumors, we performed immunohistochemical and cytokine analysis on tumors, collected 24h after the last injection (i.e., 8 and 6 days after the first injection in the CT26 and LLC models, respectively). The most striking effect of L19-mIL12 in CT26 was the marked elevation of NK cells and CD8^+^ T cells within the tumor mass. By contrast, the density of Tregs was low in all treatment groups [**Figure 4A**]. The density of NK cells and CD8^+^ T cells was also increased in LLC tumors. However, in this model, a substantial increase in Tregs was observed for all mice treated with L19-mIL12 (alone or in combination) [**Figure 4B**]. Protein extracts from tumors were analyzed with a multiplex cytokine analysis. We observed elevated levels of IFN-*γ* and MIG in the treatment group of both CT26 and LLC models [**Supplementary Figure 3**].

**Figure 4.**
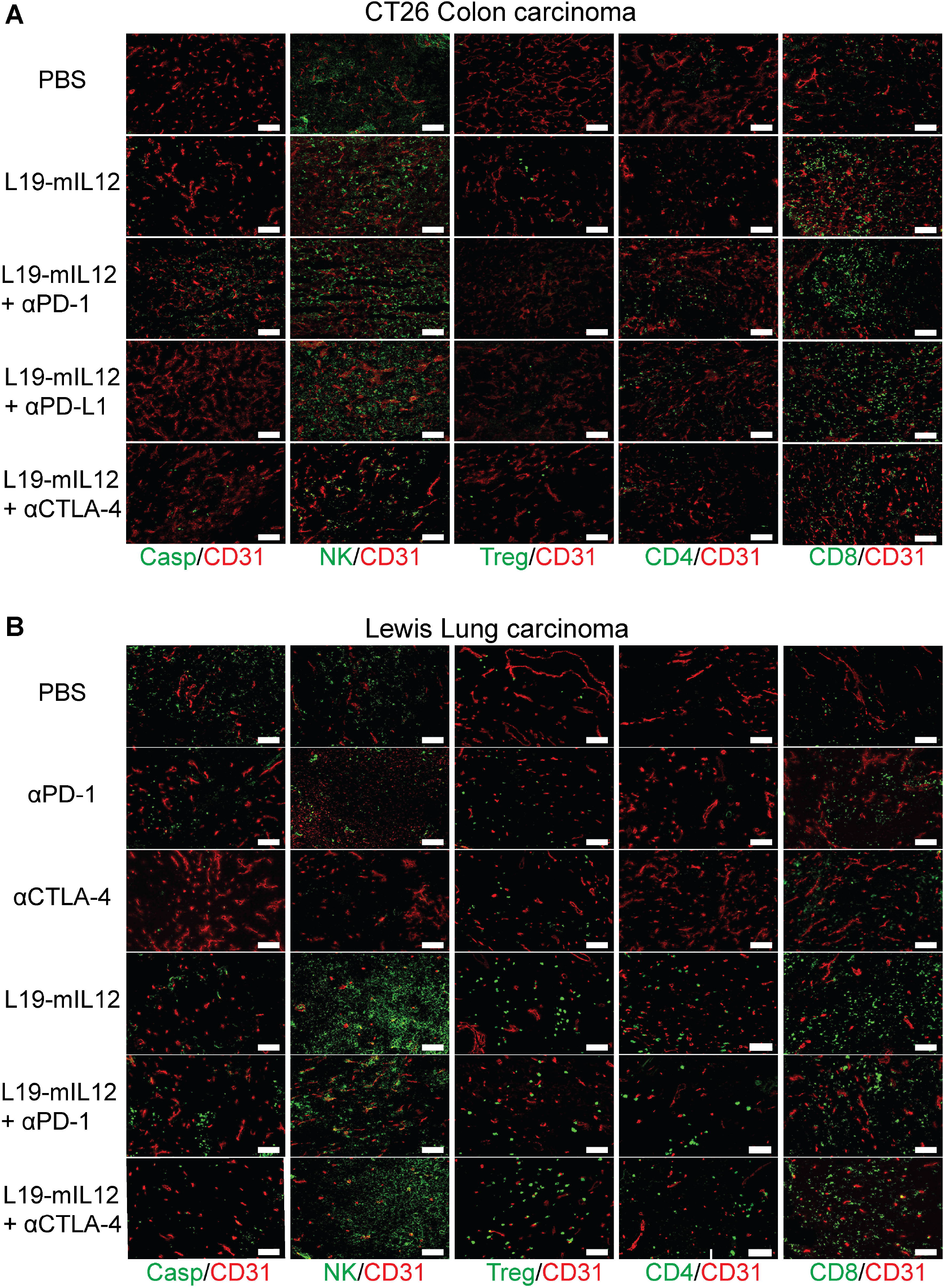
Microscopic analysis of tumor-infiltrating cells. (**A**) *Ex vivo* immunofluorescence analysis on CT26 Colon carcinoma 24h after the third injection of PBS or 12*μ*g of L19-mIL12 alone or in combination with 200*μ*g of a check-point inhibitor. Marker specific for apoptotic cells (caspase-3), NK cells (Asialo GM1), Tregs (Foxp3), CD4^+^ T cells (CD4), CD8^+^ T cells (CD8) were used (green). Blood vessels were stained with an anti-CD31 antibody (red). 10x magnification; scale bars=100*μ*m. (**B**) *Ex vivo* immunofluorescence analysis on LLC lesions 24h after the second injection of PBS, 200*μ*g αPD-1, 200 *μ*g *α*CTLA-4 or 12 *μ*g of L19-mIL12 alone or in combination with 200 *μ*g of a check-point inhibitor. The same cell markers as in (A) were used in this analysis. 10x magnification; scale bars=100*μ*m.

### Phenotype analysis of leukocytes of mice bearing CT26 carcinoma lesions

We performed a FACS characterization of leukocytes isolated from the CT26 tumor mass or from tumor-draining lymph nodes of BALB/c mice (24h after the first or third injection). Specifically, we studied features of CD4^+^ and CD8^+^ T cells, as well as a subset of these lymphocytes which are specific to AH1 (a retroviral antigen which drives anti-tumor responses in BALB/c mice) ^41–43^ [**Figure 5**]. The proportion of CD8^+^ T cells specific to AH1 was low (approximately 1%) in tumor-draining lymph nodes but increased over time both for saline and L19-mIL12 treatment groups. By contrast, 41.3% and 54.7% of the intratumoral CD8^+^ T cells were specific to AH1 after 1 and 3 injections of L19-mIL12, respectively [**Figure 5A**]. This observation suggests an active role of AH1-specific lymphocytes within the tumor mass, whose number is slightly increased by the targeted delivery of IL12 compared to the saline treatment. In tumor draining lymph nodes both CD4^+^ and CD8^+^ T lymphocytes had a naïve phenotype (i.e., CD62L^+^ CD44^−^), while the CD8^+^ AH1-specific subset showed effector memory features (i.e., CD62L^−^ CD44^+^) suggesting a specific anticancer role for this latter population. [**Figure 5B**]. These findings were paralleled by the observation of an exhausted phenotype (in terms of PD-1 expression) for CD8^+^ AH1-specific T cells, with a higher proportion of proliferating cells (Ki67 positive) in relation to the total CD8^+^ T lymphocytes [**Figure 5C**]. Analysis of the tumor bed revealed a predominantly effector memory phenotype for both CD4^+^ and CD8^+^ T cells (i.e., CD62L^−^ CD44^+^), whose frequency was not affected by the treatment [**Figure 5D**]. Moreover, intratumoral infiltrating T cells expressed high levels of immunoregulatory receptors, such as PD-1 and Tim-3, whose densities were not substantially modified by the administration of targeted IL12 [**Figure 5E**]. An analysis of vital cells within the tumor mass only revealed a significant increase in tumor cell mortality after the third injection of L19-mIL12 [**Figure 5F**], in keeping with the progressive decrease of the tumor size [**Figure 3**]. The immunohistochemical findings of **Figure 4A** on Tregs were confirmed by FACS analysis, with no substantial change in the proportion of CD4^+^ Foxp3^+^ T cells [**Figure 5G**].

**Figure 5.**
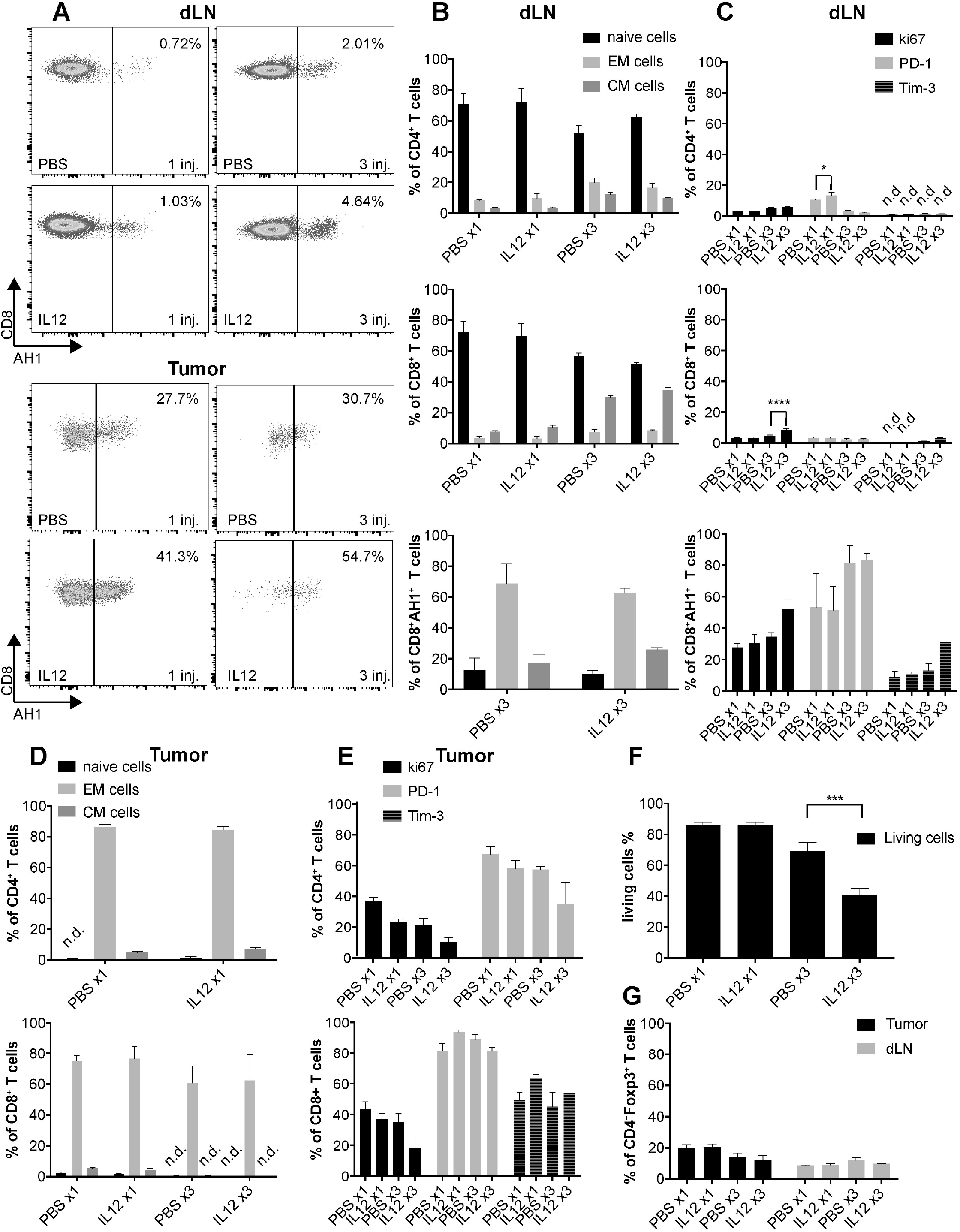
Phenotype analysis of CD4^+^, CD8^+^, CD8^+^ AH1-specific T cells in tumor draining lymphnodes (dLN) and in the tumor of CT26-bearing mice. (**A**) Frequency of AH1-specific T cells among total CD8^+^ T cells measured after 1 and 3 injections of PBS or L19-mIL12 in tumor dLN and in the tumor (n=4 mice per group). (**B**) Bar plots showing the percentage of naïve (EM; CD62L^+^ CD44^−^), effector memory (EM; CD62L^−^ CD44^+^), and central memory (CM; CD62L^+^ CD44^+^) cells among CD4^+^ T cells, CD8^+^ T cells and CD8^+^ AH1-specific T cells in tumor dLN (n=4 mice per group). (**C**) Bar plot showing the expression of immunosuppressive receptors PD-1, Tim-3 and of the proliferation marker ki67 among CD4^+^ T cells, CD8^+^ T cells and CD8^+^ AH1-specific T cells in tumor dLN (n=4 mice per group). (**D**) Bar plots showing the percentage of naïve, effector memory and central memory cells among CD4^+^ T cells, CD8^+^ T cells and CD8^+^ AH1-specific T cells in the tumor (n=4 mice per group). (**E**) Bar plot showing the expression of immunosuppressive receptors PD-1, Tim-3 and of the proliferation marker ki67 among CD4^+^ T cells, CD8^+^ T cells and CD8^+^ AH1-specific T cells in the tumor (n=4 mice per group). (**F**) Bar plot analysis showing the percentage of living tumor cells after 1 and 3 injections of PBS or L19-mIL12 (n=4 mice per group). (**G**) Bar plots representing the percentage of CD4^+^ Foxp3^+^ T cells among total CD4^+^ T cells in tumor dLN and in the tumor after 1 and 3 injections of PBS or L19-mIL12 (n=4 mice per group). Statistical differences were assessed between mice receiving the same amount of injections of PBS and L19-mIL12. *, p<0.05; ***, p<0.001; ****, p<0.0001 (regular two-way ANOVA test with Bonferroni post-test). Data represent mean (±SEM). n.d.=non-detectable.

### In vivo depletion experiment

In order to functionally confirm the contribution of various lymphocyte populations to the anti-tumor activity, we performed therapy studies in both CT26 and LLC models, following a depletion of NK cells and T cell subsets. CD8^+^ T cells and NK cells were crucially important for anti-cancer immunity in both tumor types. However, the role of CD4^+^ T cells in the LLC model was dispensable and a depletion of Tregs, mediated by an anti-CD25 antibody, did not potentiate the action of L19-mIL12 [**Figure 6**].

**Figure 6.**
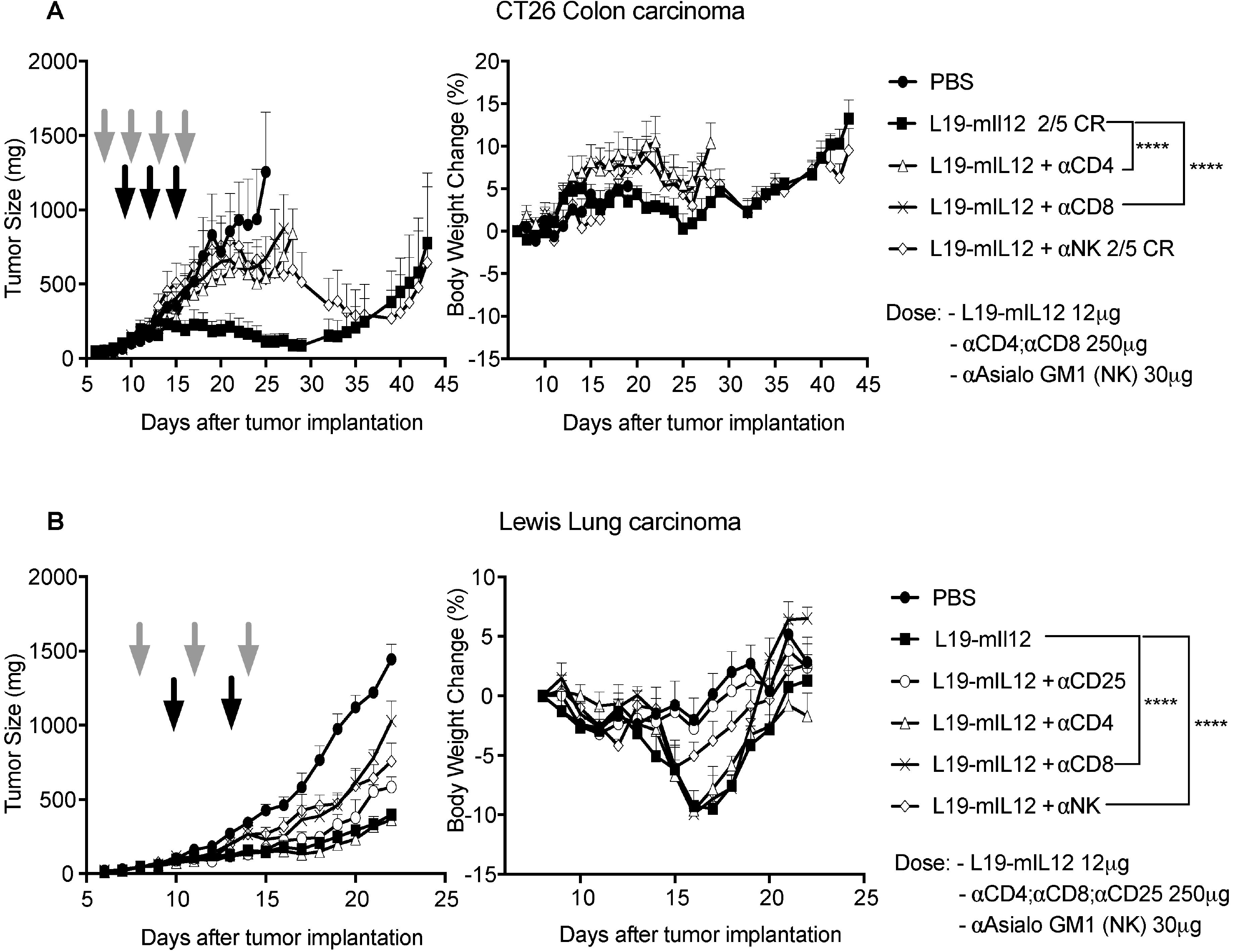
*In vivo* depletion experiment. (**A**) CT26 Colon carcinoma bearing mice received 3 intravenous injections of L19-mIL12 on days 9, 12 and 15. CD4^+^ and CD8^+^ T cell depleting antibodies were injected intraperitoneally on days 7, 10, 13 and 16 whereas NK cell depleting antibody on days 7, 10 and 13. Two undepleted groups treated either with saline or with L19-mIL12 was included as negative and positive controls respectively. n=5 mice per group. (**B**) LLC bearing mice received 2 intravenous injections of L19-mIL12 on days 10 and 13. CD25^+^, CD4^+^, CD8^+^ and NK^+^ cell depleting antibodies were injected intraperitoneally on days 8, 11 and 14. Two undepleted groups treated either with saline or with L19-mIL12 were included as negative and positive controls, respectively. Black and grey arrows indicate the injections of L19-mIL12 and the different depleting antibodies, respectively. Statistical differences were assessed between mice receiving L19-mIL12 and mice receiving L19-mIL12 plus a depleting antibody. ****, p<0.0001 (regular two-way ANOVA test with Bonferroni post-test). Data represents tumor volume ±SEM. n=4-5 mice per group. CR=Complete Response.

## DISCUSSION

We have described the production of a novel fusion protein (L19-mIL12), which selectively homes to the tumor extracellular matrix, achieving high local concentrations of murine inter-leukin-12 at the tumor site. The fusion protein was potently active against murine CT26 and WEHI-164 tumors in the immunocompetent setting, and could further be potentiated by combination with CTLA-4 or PD-1 blockade. Anti-tumor activity was also observed against LLC tumors, but in this model cancer cures could not be achieved. A possible explanation for this observation is a lower ability of L19-based immunocytokines to efficiently accumulate into LLC lesions [**Supplementary Figure 7**].

Recombinant human interleukin-12 has been used in patients, but toxicity was observed at doses as low as 1 μg/Kg body weight ^12, 44^. Objective anti-cancer responses were observed in patients with cutaneous T-cell lymphoma (56%) ^45^, non-Hodgkin’s lymphoma (21%) ^46^ and AIDS-related Kaposi’s sarcoma (50-71%) ^47^. The targeted delivery of IL12 to the tumor site mediates a substantial improvement of the therapeutic index in immunocompetent rodent models of cancer ^48^. It is thus conceivable, that a targeted approach using fully-human IL12-based products may provide a benefit to certain groups of cancer patients. Two antibody-IL12 fusion proteins have recently been studied in phase I clinical trials ^16, 17^ to determine tolerability, safety and Maximal Tolerated Dose (MTD) in patients with various advanced solid tumors. The activity of NHS-IL12 and BC1-IL12 was strongly associated with an increased serum concentration of proteins physiologically induced by IL12 (e.g., IFN-*γ* and IP10). Interestingly, patients in both trials experienced a diminished rise of circulating IFN-*γ* and IP10 upon multiple dosages, which may indicate counter regulating mechanisms against IL12 fostered by the patient’s own immune system. The toxicological profile, as evidenced by elevated levels of transaminases and decreased White Blood Cells count, was similar for both therapeutics ^16, 17^.

In mice, L19-mIL12 was very well tolerated at a dose of 12μg, which is equivalent to the human dose of 2mg, under consideration of the body surface scaling factor ^16, 17^. The lack of toxicity was corroborated by the absence of weight loss [Figure 3B **and** C] and by the histopathological analysis of organs by necropsy **[Supplementary Figure 2]**. Cancer cures for CT26 carcinomas correlated with a local increase of CD8^+^ T cells and NK cells in tumors, following administration of the antibody-IL12 fusion protein. As expected, the majority of T cells in the neoplastic lesions were different from those in secondary lymphoid organs, with a predominance of AH1-specific CD8^+^ T cells ^41–43^ [**Figure 5**]. Despite the fact that these cells displayed an exhausted phenotype in terms of PD-1 expression, they still contributed to the therapy outcome, as indicated by the lymphocyte depletion studies. High levels of PD-1 and Tim-3 were found also on AH1-specific CD8^+^ T cells in the spleen of BALB/c mice that had successfully rejected a tumor rechallenge after a first cancer cure [**Supplementary Figure 8**].

It is becoming increasingly clear that aberrantly expressed tumor-rejection antigens (including peptides corresponding to genes of retroviral origin, which have been integrated in the genome at some stage during evolution) may be relevant for anti-cancer immune surveillance in mice as well as in humans ^49^. Experimental evidence arises not only from mass spectrometry-based analysis of peptide presentation on MHC class I molecules ^43, 49^, but also from the results of therapeutic vaccination studies with peptides corresponding to aberrantly expressed antigens ^49, 50^. When considering human tumors, Miller et al. have reported that polyomavirus-specific T cells may represent up to 21% of the T cell specificities within the tumor mass of Merkel Cell Carcinoma (MCC) ^51^. Besides virally-driven tumors (e.g., MCC and HPV-derived malignancies), twenty potentially immunogenic endogenous retroviruses have recently been reported in clear-cell renal cell carcinoma patients ^52^. These findings may explain why renal tumors are immunogenic (i.e., respond well to PD-1 blockade ^53, 54^ or to interleukin-2 ^55^), in spite of not having a particularly high mutation prevalence ^56^.

Interestingly, L19-mIL12 triggered a local intratumoral increase of Tregs in LLC lesions, but not in CT26 tumors. Both cancer types expressed comparable amounts of the EDB domain of fibronectin, the target of L19. Depletion of CD25-positive T cells did not potentiate the anti-cancer activity of L19-mIL12 [**Figure 6**]. It is possible, however, that tumor-targeting antibody-IL12 fusions may display their therapeutic effect, at least in part, by modulating Treg fragility at the site of disease ^57^. The targeted delivery of interleukin-12 to the neoplastic mass leads to high levels of IFN-*γ* within the tumor mass ^28^, a cytokine which acts as key modulator of Treg functionality ^57^.

Therapy studies with lymphocyte depletion have shown that, in addition to CD8^+^ T cells, also NK cells play an important role in the tumor rejection process in mice, which had received L19mIL12 [**Figure 6**]. Stress proteins (e.g., H60a, b, c; Rae-1α-ε and Mult1) upregulated on the surface of tumor cells can be efficiently recognized by activating receptors on NK cells, such as NKG2D ^58, 59^.

Various formats of antibody-IL12 fusions have been considered over the year ^28–30, 60, 61^. The format of L19-mIL12, similar to the one previously published by our group for the F8 antibody ^30^ appears to have advantages compared to other formats, both in terms of ease of production and of biochemical quality. L19-mIL12 is a single polypeptide, which avidly binds to the cognate antigen as it contains two antibody moieties. Other formats, featuring interleukin-12 as disulfide-bonded heterodimeric structures, suffered from the undesired formation of homodimeric products and from challenges in fermentation procedures ^29, 60^. L19-mIL12 exhibited potent single agent activity, which was enhanced by combination with immune checkpoint inhibitors. In keeping with our findings, the combination of NHS-IL12 (an IL12 fusion based on an antibody directed against DNA released from necrotic tumor cells in full IgG format) with PD-L1 blockade had also revealed a synergistic activity ^62^. However, no biodistribution studies with IL12 fusions in IgG format have been published so far and it is therefore difficult, at this stage, to assess whether these molecules have equivalent activity. In principle, the tandem diabody format (used in this work) should promote an easier extravasation and tumor targeting process, while rapidly clearing from circulation. Indeed, the biodistribution results of **Figure 2** reveal favorable tumor:organ ratios, 24h after i.v. administration. A rapidly-clearing antibody-cytokine fusion should also be easier to administer to patients, as infusions could be stopped upon onset of side-effects, which usually happens at peak concentrations of the cytokine moiety in blood ^18, 20, 63^. Collectively, the data presented in this article provide a rationale for the clinical investigation of fully human analogues of L19-mIL12, which feature human interleukin-12 as therapeutic payload.

## ACKNOWLEDGEMENTS

This project has received funding from the European Research Council under the Grant Agreement No 670603, ETH Zürich, the Swiss National Science Foundation (project number: 310030_182003/1), the Swiss Federal Commission for Technology and Innovation (Grant number: 17072.1) and the “Stiftung zur Krebsbekämpfung”. Help from Diana Tintor for a critical reading of the manuscript.

