## Supplementary Figure Legends for "The antibody-based delivery of interleukin-12 to solid tumors boosts NK and CD8^+^ T cell activity and synergizes with immune check-point inhibitors"

**Supplementary Figures Legends**

**Supplementary Fig.S1.**

**(A**) Gene sequence and (**B**) full amino acid sequence of L19-mIL12.

**Supplementary Fig.S2**.

Histological assessment of HE-stained tissue sections in healthy BALB/c mice treated with Saline (n=1) or with 3 injections of L19-mIL12 (n=3). (**A**) Mild multifocal inflammatory cell infiltrate in the Liver of a mouse treated with L19-mIL12 (central and right figures) compared to control (left figure). H=Hematopoietic cell precursors; M=Macrophages; PP=Periportal Vein. Scale bar: left and central figure 100μm; right figure=50μm. (**B**) Spleen of mice treated with L19-mIL12 showed moderately increased extramedullary haematopoiesis (arrows) compared to control. WP=White Pulp; RP=Red Pulp. Scale bar=1mm. (**C**) Lymph nodes of mice in the treatment group showed a mild increase in macrophages in the paracortex and medullary sinuses (arrows). Scale bar=100μm. (**D**) Mild myeloid hyperplasia (arrows) in the bone marrow of mice receiving L19-mIL12. Scale bar=50μm. (**E**) Scattered apoptotic tubular epithelial cells (A) in the kidney of mice treated with L19-mIL12. Scale bar=100m. (**F**) Minimal myofibre degeneration (arrows) with inflammation in the quadriceps (this lesion was observed also in muscles adjacent to the spinal column, muscles of the head and tongue; the heart was not affected) of mice treated with L19-mIL12. Scale bar=250μm.

**Supplementary Fig.S3**.

Cytokine analysis of (**A**) CT26 Colon carcinoma and (**B**) Lewis Lung carcinoma extracts using multiplexed particle-based flow cytometry assay. Statistical differences were assessed between treatment and control groups. **, p<0.01; ****, p<0.0001 (regular two-way ANOVA test with Bonferroni post-test). (n=2 mice per group).

**Supplementary Fig.S4**.

(**A**) *Ex vivo* immunofluorescence analysis on CT26 Colon carcinoma 24h after the third injection of PBS or 12$\mu$g of L19-mIL12 alone or in combination with 200$\mu$g of a check-point inhibitor. Marker specific for NK cells (NKp46), was used (green). Blood vessels were stained with an anti-CD31 antibody (red). 10x magnification; scale bars=100$\mu$m. (**B**) *Ex vivo* immunofluorescence analysis on LLC lesions 24h after the second injection of PBS, 200$\mu$g $\alpha$PD-1, 200$\mu$g $\alpha$CTLA-4 or 12$\mu$g of L19-mIL12 alone or in combination with 200$\mu$g of a check-point inhibitor. The same cell marker as in (A) was used in this analysis. 10x magnification; scale bars=100$\mu$m.

**Supplementary Fig.S5**.

Therapy study in BALB/c mice bearing CT26 lesions. Treatment started when tumors reached 100mm^3^. Mice received 200µg of $\alpha$PD-1 or$\alpha$PD-L1 or $\alpha$CTLA-4 antibody, 3 times every 72h (n=4 mice per group).

**Supplementary Fig.S6**.

Quantification of the leukocyte density depicted in **Figure 4** using ImageJ software.

**Supplementary Fig.S7**.

Quantitative biodistribution experiment of 30µg radioiodinated L19-IL2 24h after intraveneous injection in immunocompetent Lewis Lung carcinoma bearing mice (C57BL/6). Results are expressed as percentage of injected dose per gram of tissue (%ID/g $\pm$SEM; n=5 mice).

**Supplementary Fig.S8**.

Analysis 8 days after a tumor rechallenge of AH1-specific CD8^+^ T cells in the spleen of BALB/c mice cured by the treatment of L19-mIL12 [**Figure 3B**]. All the mice (n=10) not sacrificed for this study successfully rejected the rechallenge with CT26 Colon carcinoma cells. (n=2 mice analysed in the study).
