## Supplementary Figure 1 for "The antibody-based delivery of interleukin-12 to solid tumors boosts NK and CD8^+^ T cell activity and synergizes with immune check-point inhibitors"

**Supplementary Fig.S1.**

**A**

**NheI//** Leader Sequence-mIL12 beta p40-Linker-mIL12 alpha p35- Linker- L19 -Linker –L19 **//NotI**

CTCCGCTAGCGTCGACCATGGGCTGGAGCCTGATCCTCCTGTTCCTCGTCGCTGTGGCTACAGGTgtgcacTCGatgtgggagctggagaaagacgtttatgttgtagaggtggactggactcccgatgcccctggagaaacagtgaacctcacctgtgacacgcctgaagaagatgacatcacctggacctcagaccagagacatggagtcataggctctggaaagaccctgaccatcactgtcaaagagtttctagatgctggccagtacacctgccacaaaggaggcgagactctgagccactcacatctgctgctccacaagaaggaaaatggaatttggtccactgaaattttaaaaaatttcaaaaacaagactttcctgaagtgtgaagcaccaaattactccggacggttcacgtgctcatggctggtgcaaagaaacatggacttgaagttcaacatcaagagcagtagcagttcccctgactctcgggcagtgacatgtggaatggcgtctctgtctgcagagaaggtcacactggaccaaagggactatgagaagtattcagtgtcctgccaggaggatgtcacctgcccaactgccgaggagaccctgcccattgaactggcgttggaagcacggcagcagaataaatatgagaactacagcaccagcttcttcatcagggacatcatcaaaccagacccgcccaagaacttgcagatgaGgcctttgaagaactcacaggtggaggtcagctgggagtaccctgactcctggagcactccccattcctacttctccctcaagttctttgttcgaatccagcgcaagaaagaaaagatgaaggagacagaggaggggtgtaaccagaaaggtgcgttcctcgtagagaGgacatctaccgaagtccaatgcaaaggcgggaatgtctgcgtgcaagctcaggatcgctattacaattcctcgtgcagcaagtgggcatgtgttccctgcagggtccgatccGGTGGAGGCGGTTCAGGCGGAGGTGGCTCTGGCGGTGGCGGATCGagggtcattccagtctctggacctgccaggtgtcttagccagtcccgaaacctgctgaagaccacagatgacatggtgaagacggccagagaa***aaGctT***aaacattattcctgcactgctgaagacatcgatcatgaagacatcacacgggaccaaaccagcacattgaagacctgtttaccactggaactacacaagaacgagagttgcctggctactagagagacttcttccacaacaagagggagctgcctgcccccacagaagacgtctttgatgatgaccctgtgccttggtagcatctatgaggacttgaagatgtaccagacagagttccaggccatcaacgcagcacttcagaatcacaaccatcagcagatcattctagacaagggcatgctggtggccatcgatgagctgatgcagtctctgaatcataatggcgagactctgcgccagaaacctcctgtgggagaagcagacccttacagagtgaaaatgaagctctgcatcctgcttcacgccttcagcacccgcgtcgtgaccatcaacagggtgatgggctatctgagctccgccGGTAGCGCTGATGGAgaggtgcagctgttggagtctgggggaggcttggtacagcctggggggtccctgagactctcctgtgcagcctctggattcacctttagcagtttttcgatgagctgggtccgccaggctccagggaaggggctggagtgggtctcatctattagtggtagttcgggtaccacatactacgcagactccgtgaagggccggttcaccatctccagagacaattccaagaacacgctgtatctgcaaatgaacagcctgagagccgaggacacggccgtatattactgtgcgaaaccgtttccgtattttgactactggggccagggaaccctggtcaccgtctcgagtgggtccagtggcggtgaaattgtgttgacgcagtctccaggcaccctgtctttgtctccaggggaaagagccaccctctcctgcagggccagtcagagtgttagcagcagctttttagcctggtaccagcagaaacctggccaggctcccaggctcctcatctattatgcatccagcagggccactggcatcccagacaggttcagtggcagtgggtctgggacagacttcactctcaccatcagcagactggagcctgaagattttgcagtgtattactgtcagcagacgggtcgtattccgccgacgttcggc*caagggaccaaggtggaaatcaaa*tcttcctcatcCggAagtagctcttcG***ggAtcC***tcgtccagcggcgaggtgcagctgttggagtctgggggaggcttggtacagcctggggggtccctgagactctcctgtgcagcctctggattcacctttagcagtttttcgatgagctgggtccgccaggctccagggaaggggctggagtgggtctcatctattagtggtagttcgggtaccacatactacgcagactccgtgaagggccggttcaccatctccagagacaattccaagaacacgctgtatctgcaaatgaacagcctgagagccgaggacacggccgtatattactgtgcgaaaccgtttccgtattttgactactggggccagggaaccctggtcaccgtctcgagtgggtccagtggcggtgaaattgtgttgacgcagtctccaggcaccctgtctttgtctccaggggaaagagccaccctctcctgcagggccagtcagagtgttagcagcagctttttagcctggtaccagcagaaacctggccaggctcccaggctcctcatctattatgcatccagcagggccactggcatcccagacaggttcagtggcagtgggtctgggacagacttcactctcaccatcagcagactggagcctgaagattttgcagtgtattactgtcagcagacgggtcgtattccgccgacgttcggc*caagggaccaaggtggaaatcaaa*TAGTGA**GCGGCCGC**AAAAGGAAAA

**B**

mIL12 p40-Linker-mIL12 p35- Linker- L19 -Linker –L19

MWELEKDVYVVEVDWTPDAPGETVNLTCDTPEEDDITWTSDQRHGVIGSGKTLTITVKEFLDAGQYTCHKGGETLSHSHLLLHKKENGIWSTEILKNFKNKTFLKCEAPNYSGRFTCSWLVQRNMDLKFNIKSSSSSPDSRAVTCGMASLSAEKVTLDQRDYEKYSVSCQEDVTCPTAEETLPIELALEARQQNKYENYSTSFFIRDIIKPDPPKNLQMRPLKNSQVEVSWEYPDSWSTPHSYFSLKFFVRIQRKKEKMKETEEGCNQKGAFLVERTSTEVQCKGGNVCVQAQDRYYNSSCSKWACVPCRVRSGGGGSGGGGSGGGGSRVIPVSGPARCLSQSRNLLKTTDDMVKTAREKLKHYSCTAEDIDHEDITRDQTSTLKTCLPLELHKNESCLATRETSSTTRGSCLPPQKTSLMMTLCLGSIYEDLKMYQTEFQAINAALQNHNHQQIILDKGMLVAIDELMQSLNHNGETLRQKPPVGEADPYRVKMKLCILLHAFSTRVVTINRVMGYLSSAGSADGEVQLLESGGGLVQPGGSLRLSCAASGFTFSSFSMSWVRQAPGKGLEWVSSISGSSGTTYYADSVKGRFTISRDNSKNTLYLQMNSLRAEDTAVYYCAKPFPYFDYWGQGTLVTVSSGSSGGEIVLTQSPGTLSLSPGERATLSCRASQSVSSSFLAWYQQKPGQAPRLLIYYASSRATGIPDRFSGSGSGTDFTLTISRLEPEDFAVYYCQQTGRIPPTFGQGTKVEIKSSSSGSSSSGSSSSGEVQLLESGGGLVQPGGSLRLSCAASGFTFSSFSMSWVRQAPGKGLEWVSSISGSSGTTYYADSVKGRFTISRDNSKNTLYLQMNSLRAEDTAVYYCAKPFPYFDYWGQGTLVTVSSGSSGGEIVLTQSPGTLSLSPGERATLSCRASQSVSSSFLAWYQQKPGQAPRLLIYYASSRATGIPDRFSGSGSGTDFTLTISRLEPEDFAVYYCQQTGRIPPTFGQGTKVEIK
