## Supplementary figures and images for "The antibody-based delivery of interleukin-12 to solid tumors boosts NK and CD8^+^ T cell activity and synergizes with immune check-point inhibitors"

### Supplementary Figure 3

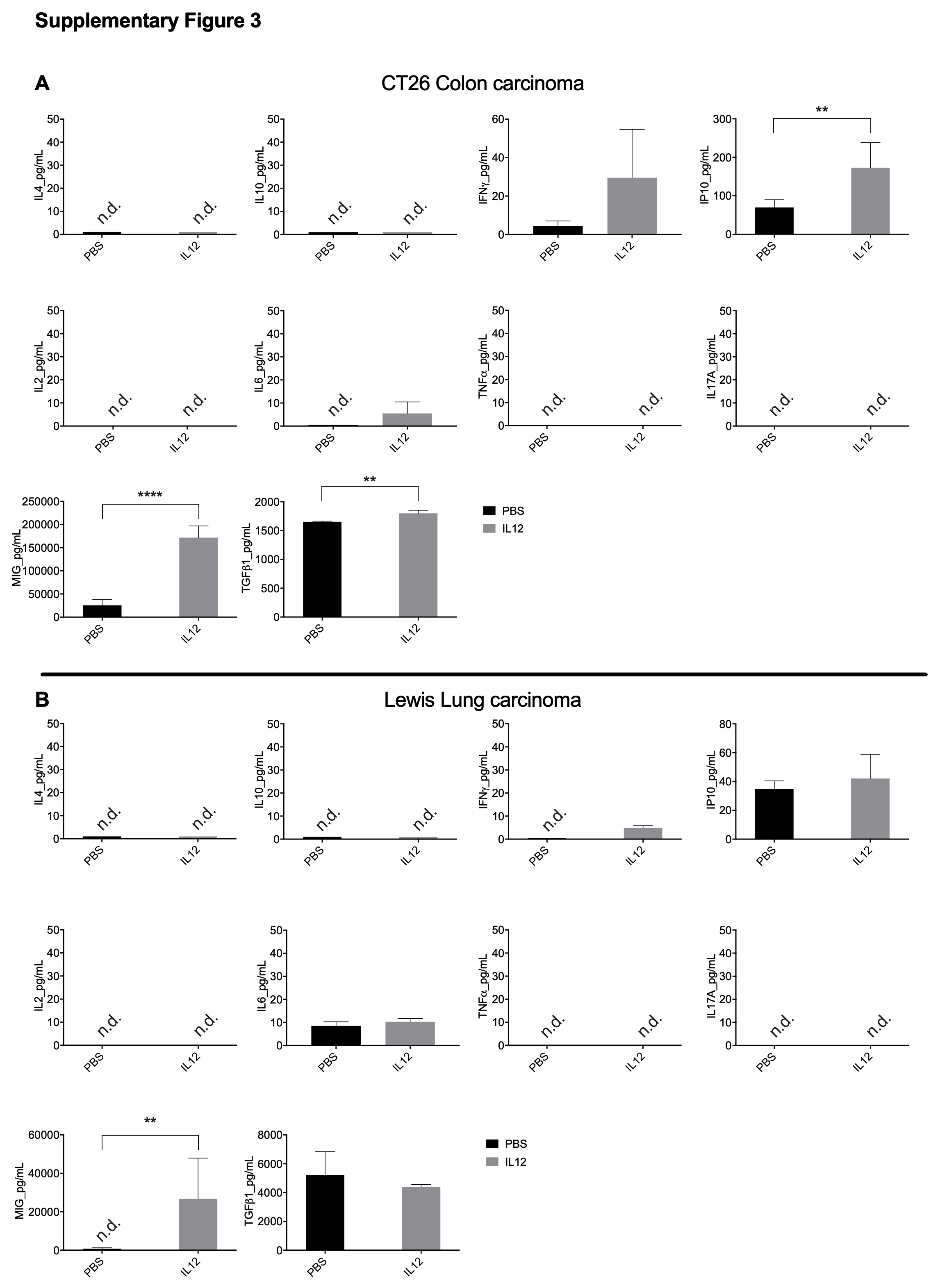

### Supplementary Figure 4

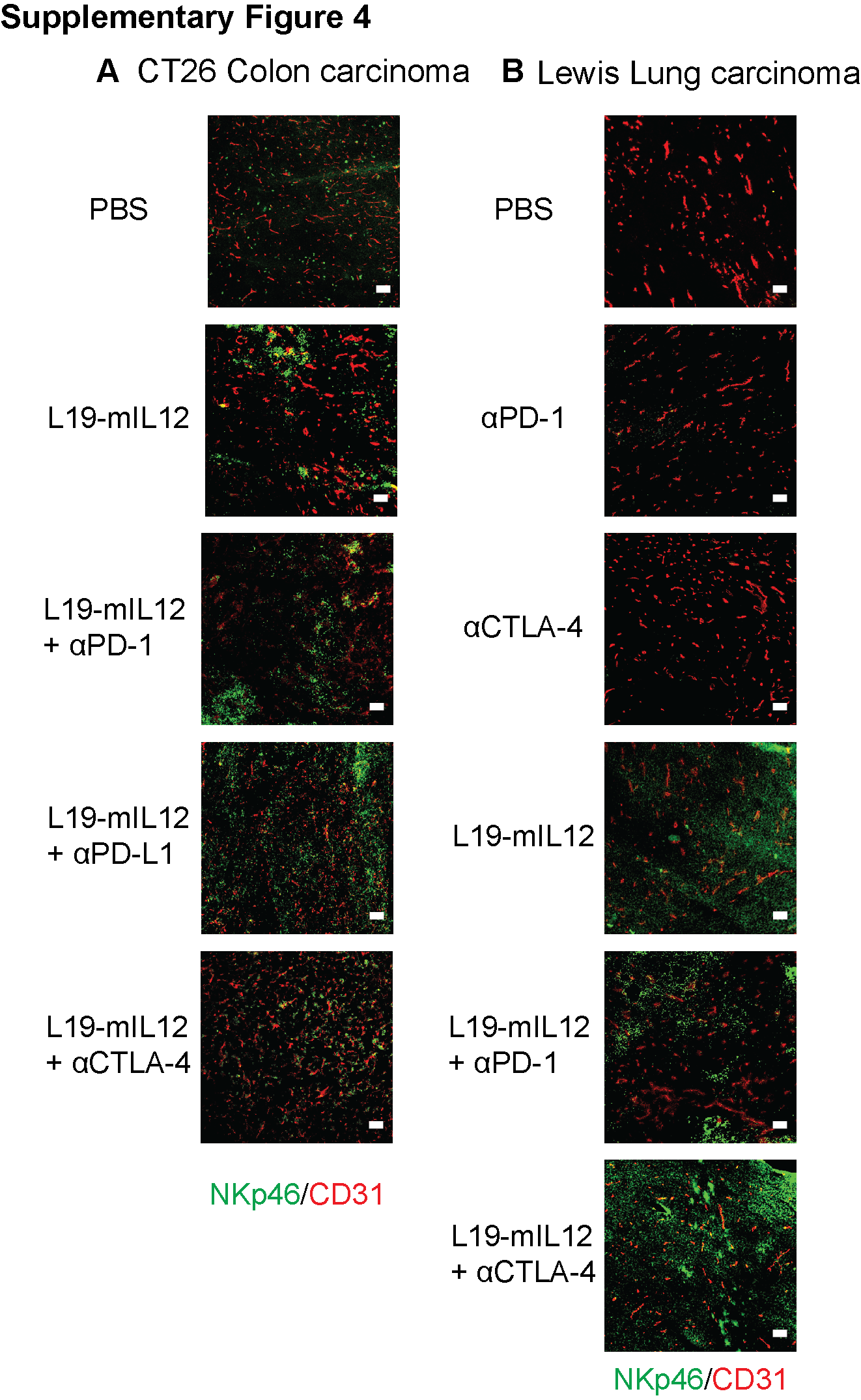

### Supplementary Figure 5

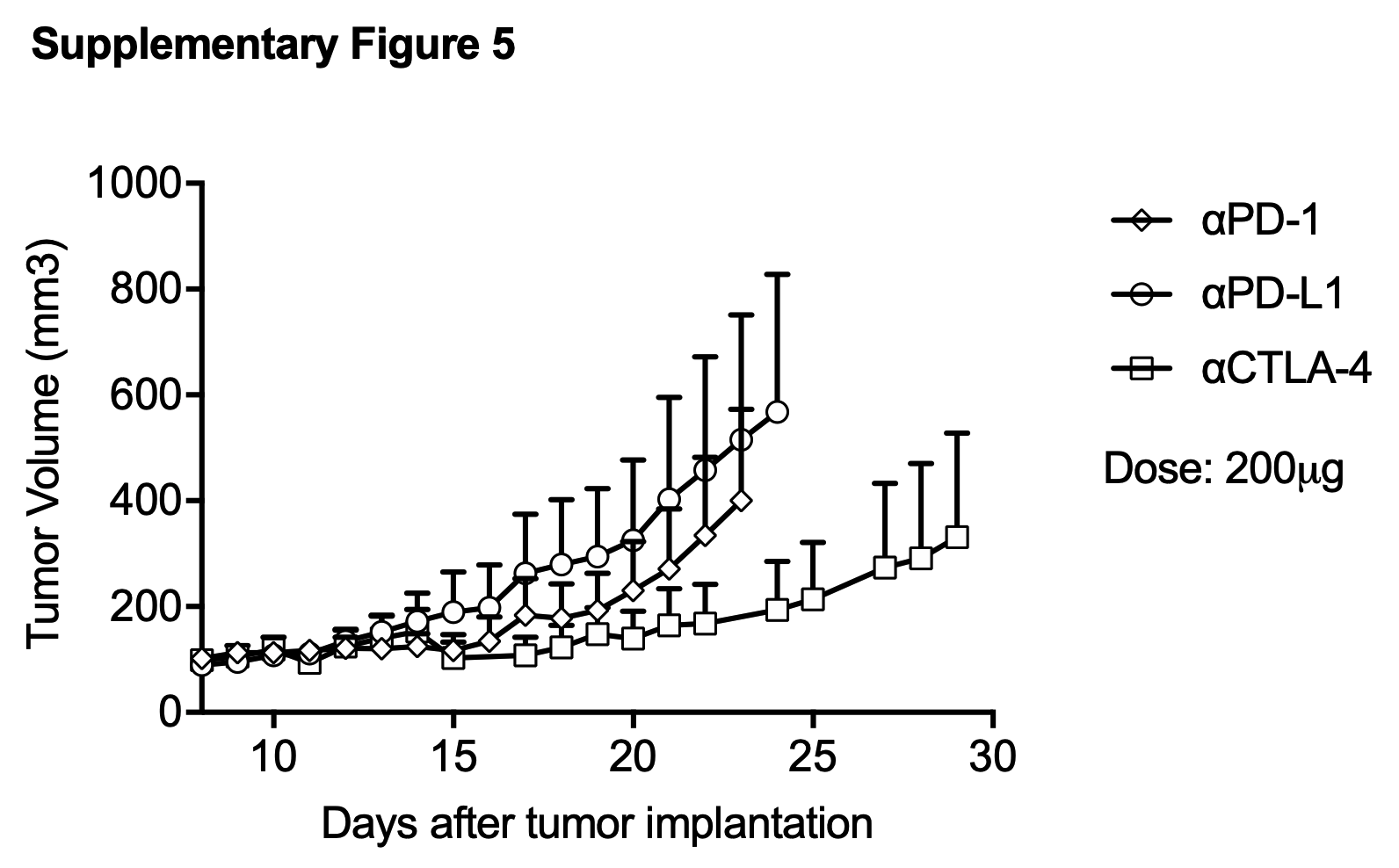

### Supplementary Figure 6

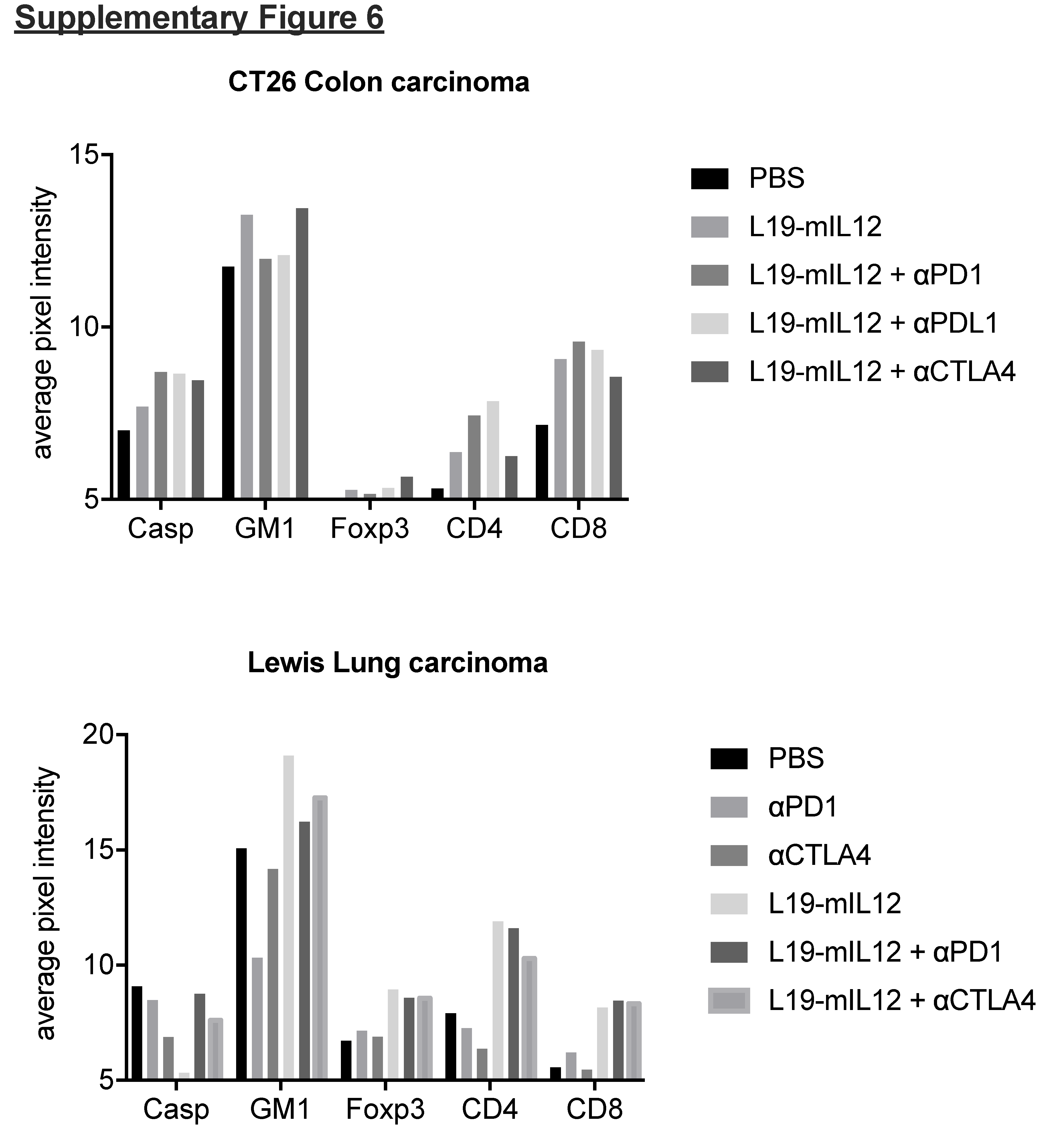

### Supplementary Figure 7

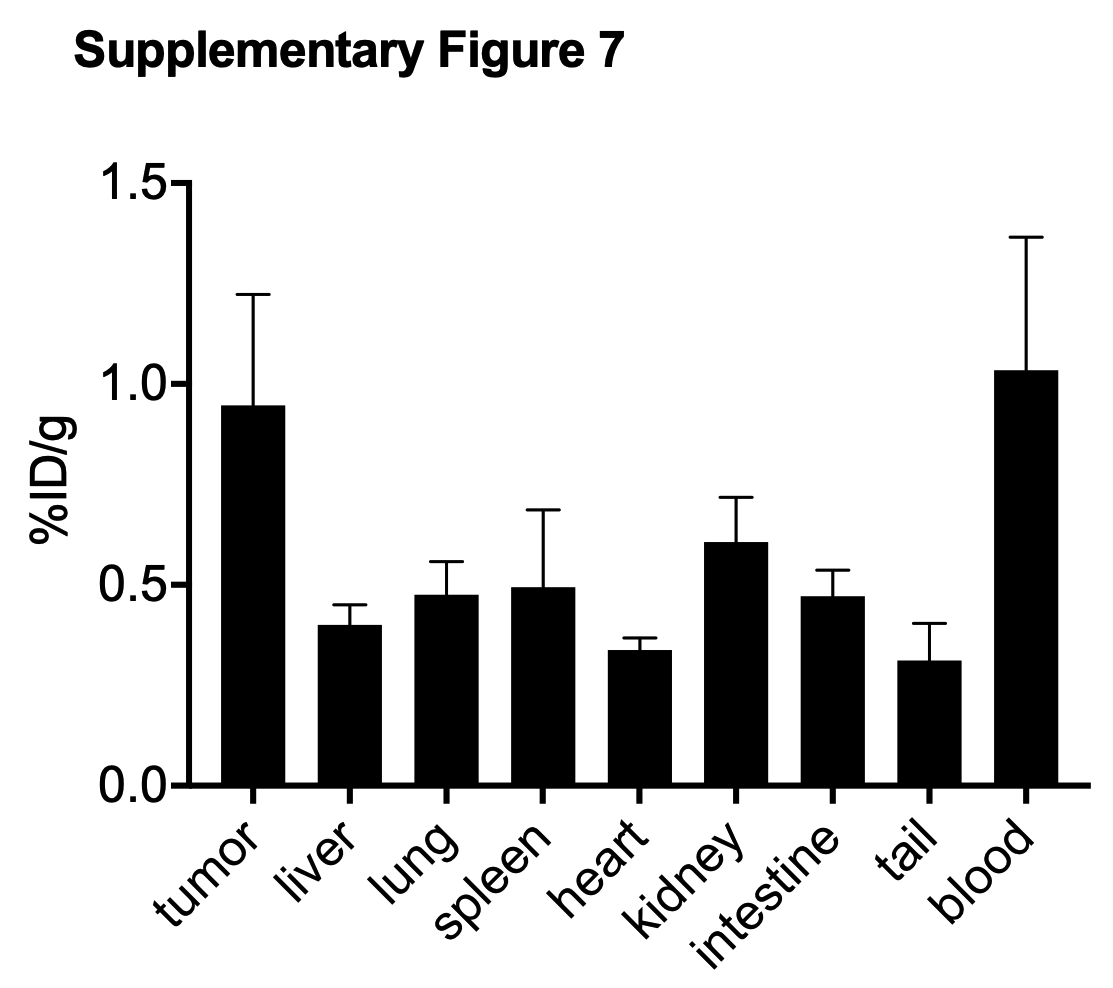

### Supplementary Figure 8

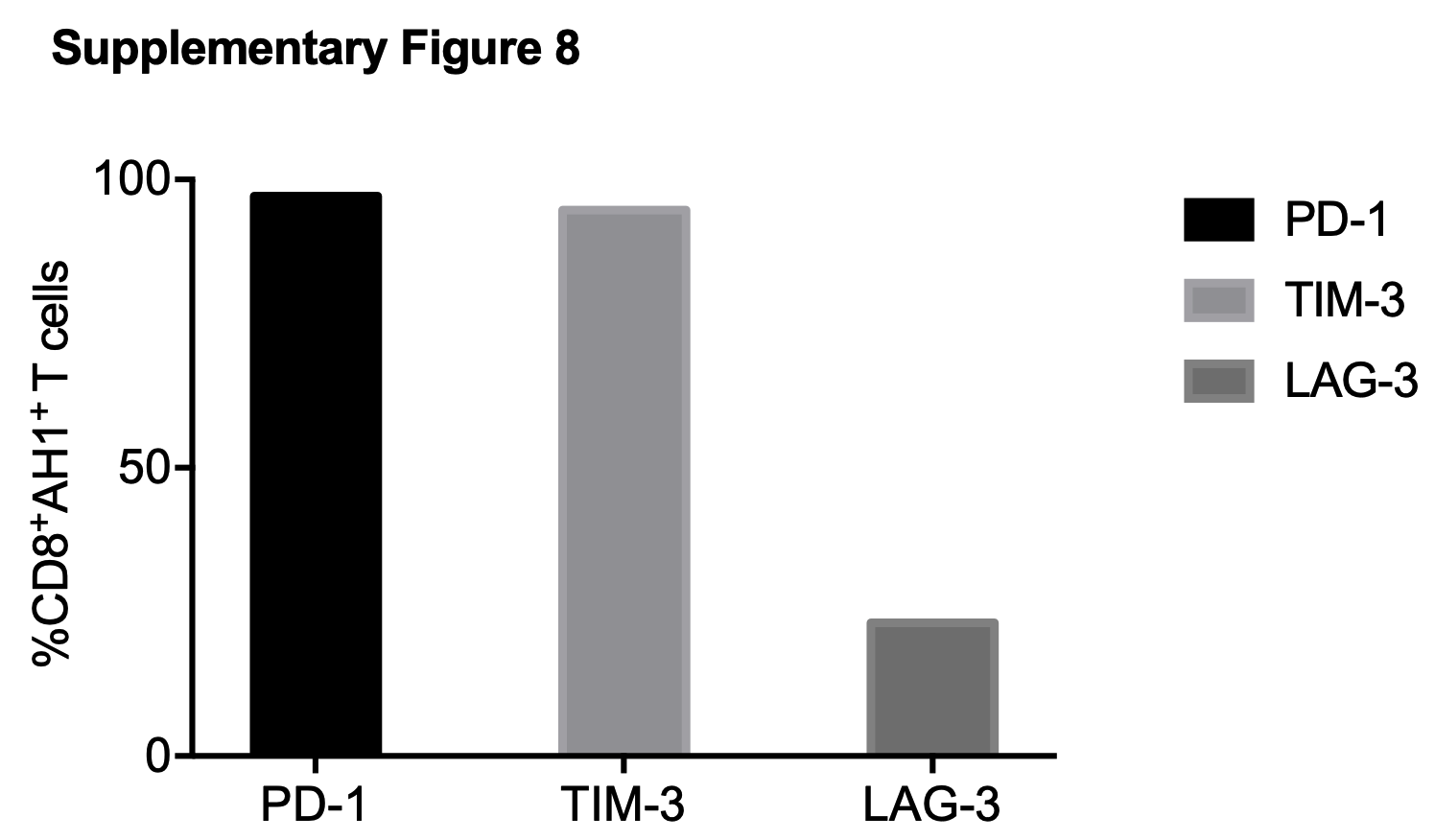
